# Models of primate ventral stream that categorize and visualize images

**DOI:** 10.1101/2020.02.21.958488

**Authors:** Elijah Christensen, Joel Zylberberg

## Abstract

An open question in systems neuroscience is which objective function (or computational “goal”) best describes the computations performed by the ventral stream (VS) of primate visual cortex. Substantial past research has suggested that object categorization could be such a goal. Recent experiments, however, showed that information about object positions, sizes, etc. is encoded with increasing explicitness along this pathway. Because that information is not necessarily needed for object categorization, this motivated us to ask whether primate VS may do more than “just” object recognition. To address that question, we trained deep neural networks, all with the same architecture, with three different objectives: a supervised object categorization objective; an unsupervised autoencoder objective; and a semi-supervised objective that combined autoencoding with categorization. We then compared the image representations learned by these models to those observed in areas V4 and IT of macaque monkeys using canonical correlation analysis (CCA). We found that the semi-supervised model provided the best match the monkey data, followed closely by the unsupervised model, and more distantly by the supervised one. These results suggest that multiple objectives – including, critically, unsupervised ones – might be essential for explaining the computations performed by primate VS.

## Introduction

The ventral stream (VS) of visual cortex begins in primary visual cortex (V1), ends in inferior temporal cortex (IT), and is essential for object recognition. Accordingly, a long-standing hypothesis in the field is that the ventral stream could be understood as mapping visual scenes onto neuronal firing patterns that represent object identity^1–5^. Supporting that assertion, deep convolutional neural networks (DCNN’s) trained to categorize objects in natural images develop intermediate representations that resemble those in primate VS^2,6–8^. At the same time, VS and other visual areas are also engaged during visualization of both previously encountered and novel scenes^9,10^, suggesting that the VS can *generate* visual scenes in addition to identifying objects within those scenes. Furthermore, non-categorical information, about object positions^11^, sizes, etc. is also represented with increasing explicitness in late VS areas V4 and IT^12^. This non-categorical information is not necessarily needed for object recognition tasks, although interestingly, deep convolutional neural networks (CNNs) recapitulated this trend of increasingly explicit category-orthogonal representations with increasing depth^11^. Nevertheless these recent findings motivated us to reconsider the long-standing question: What computational objective best explains VS physiology^13–15^?

To address this question, we pursued a recently-popularized approach^2,7,8,12,14,15^ and trained deep neural networks to perform one of three different tasks, each of which corresponds to a different computational objective. We trained the networks to either: a) recognize objects; b) form compressed image representations that suffice for reconstructing the input image; or c) recognizing objects while *also* retaining enough information about the input image to allow its reconstruction. We then compared these trained neural networks’ responses to image stimuli to responses observed in neurophysiology experiments wherein monkeys saw the same images that were input to the models, to see which tasks yielded models that best matched the neural data. We used the same architecture for all of these networks, ensuring that any differences in how well the models recapitulate the neural data can be attributed to their objective function, and not to architecture differences. Our main finding is that networks trained with objective (c) provided the closest match for both areas V4 and IT of the monkey, closely followed by ones trained with objective (b), and more distantly followed by the networks trained on the pure object recognition objective (a). This suggests that a full understanding of visual ventral stream computations might require considerations beyond object recognition, and that scene reconstruction is a promising candidate for the “other” computations occurring within the VS. Notably, other work^14–18,^, including two concurrent studies^14,15^, has asked whether unsupervised image processing models can describe primate VS function. We discuss our findings in the context of these concurrent studies in the Discussion.

## Results

### Computational Models

To identify the degree to which different computational objectives describe ventral stream physiology, we optimized deep convolutional neural network (CNN) models for different objectives, and compared them to neural recordings from the primate ventral stream. Each computational model was constructed out of a series of layers of artificial neurons, connected sequentially. The first layer takes as input an image ***x*** and at the final layer outputs a set of neuronal activities that represent the visual scene input (Fig 1B), including object identity. We refer to this output as the *latent representation*. The input images, ***x***, consisted of images of clothing articles superimposed over natural image backgrounds (see Methods). Each image used a single clothing article rendered in a randomly chosen position and orientation, and placed over a natural image background (Fig. 1A).

**Fig. 1:**
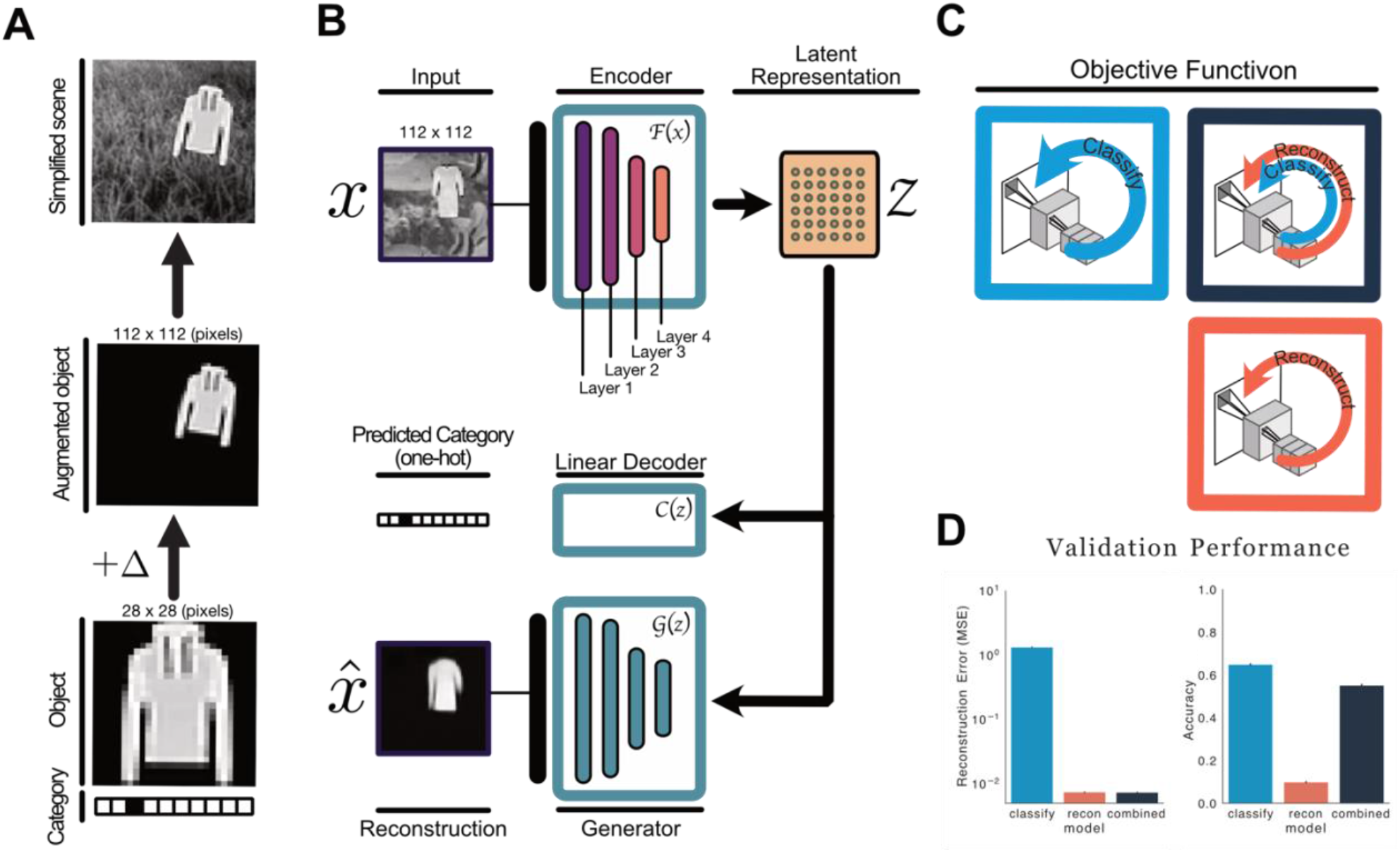
Overview. **A)** We constructed images of clothing items superimposed over natural image backgrounds at random eccentricities and orientations. **B)** We model the ventral stream as an encoder whose objective is to map input image (x) onto more abstract “latent” representations (*z*). In our models this latent space contains 500 artificial neurons. The latent layer (*z*) is densely connected whereas the preceding layers were all convolutional (see Methods). The generator network (G) uses these latent representations (*z*) as input to reconstruct the object at the correct location within the scene. A separate linear decoder attempts to determine the object identity from the activities of the units in *z*. **C)** We trained these neural networks on one of three tasks: object categorization (“classify”), object reconstruction (“reconstruct”), or object categorization with concurrent image reconstruction (“combined”). **D)** Object categorization and reconstruction performance of the three networks after they were trained, assessed on held-out images (i.e., ones not used in training the networks).

The models each had a total of four layers of processing between their inputs and these latent representations. The visual inputs to the model had normalized luminance values, mimicking the normalization observed at LGN^19^. The connectivity between neurons in each layer (and the artificial neurons’ biases) were optimized within each model, to achieve the specified objective (see Methods). We repeated this process for three different objectives, yielding three different types of models. The first type of model was optimized strictly for object recognition: the optimization maximized the ability of a linear decoder to determine the identity of the clothing object in the visual scene from the latent representation. (This mirrors the observation that neural activities in area IT can be linearly decoded to recover object identity^12^). We refer to this network as the “classify” network. The second type of model was optimized for the ability of a decoder network to reconstruct the object from the latent representation. We refer to this autoencoder as the “reconstruct” model. Finally, we considered a model whose objective during training is the sum of the “classify” objective and the “reconstruct” one: the optimization simultaneously maximized this network’s ability to perform both tasks, and we refer to it as the “combined” model. This combined model is a semi-supervised autoencoder, the construction of which was motivated by previous work in machine learning^20^.

In all cases, the models were optimized via backpropagation using sets of images containing randomly sampled objects, until their object classification performance saturated on a set of held-out validation images. Reasonable performance on the categorization task was obtained the “classify” and “combined” models (Fig 1D); as expected, the “reconstruct” model had very poor classification performance. Similarly, we assessed the ability of an optimized generator network to decode the latent state activations to reconstruct the input images. After training, both the “combined” model, and the “reconstruct” model, had relatively low reconstruction errors, whereas the “classify” model, had much higher reconstruction error. Thus, we created neural networks that could either classify image contents but not reconstruct the images themselves (“classify”), reconstruct but not classify (“reconstruct”), or do both tasks with reasonable efficacy (“combined”).

Having developed models optimized for these different objectives, we could evaluate how well each model matched observations from primate VS, and use that comparison to determine which computational objective provides the best description of primate VS.

### Electrophysiology Comparisons

To compare our neural network models to ventral stream physiology, we used the experimental data from a previously-published study^12,21^ (see Methods and Refs. 12,21 for details). These data consisted of electrode array recordings from areas V4 and IT of monkeys that were viewing images of objects superimposed over natural image backgrounds, at different locations and orientations. Many neurons in each area were simultaneously observed in these experiments.

First, we asked how well each layer within each neural network model matched the primate VS data. To achieve this goal, we input into our models the same images that were shown to the monkeys in the physiology experiments. We then extracted the activations of the artificial neurons at each layer of our computational models, and we used Canonical Correlation Analysis (CCA)^22,23^ to compare those artificial neurons’ activations to those recorded in monkey V4 and IT (See Methods). In brief, CCA assesses the degree to which weighted sums of our neural network unit activations correlate with weighted sum of the neuron firing rates observed in the monkey experiments. It can thus test for similarity in how the images are represented by the neural networks, and the monkey, without requiring us to assign each neural network unit to a specific neuron in the monkey experiments. Similar to regular correlation analysis, CCA correlations of 0 indicate no relation between the neural network and monkey visual representations, while a value of 1 indicate perfect similarity. We extracted the canonical correlations for the first 10 CCA components, and averaged their values (Fig. 2).

**Fig. 2:**
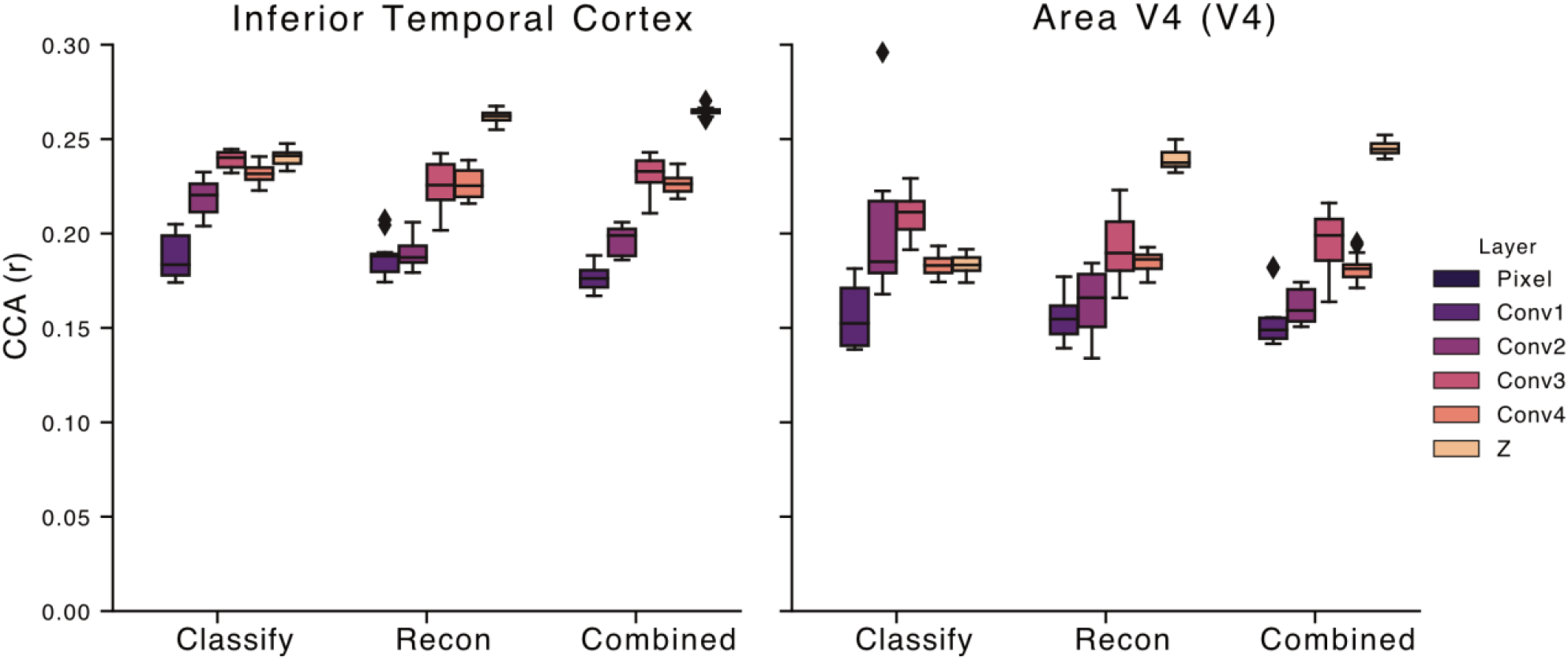
Canonical Correlation Analysis. We used Canonical Correlation Analysis (CCA) to quantify how similar the responses in the layers of each model were to primate electrophysiology data in both inferior temporal cortex (IT) and visual area V4 (V4). We used random draws of 250 unit activations in each layer of the fully trained convolutional models optimized under the “classify” objective (categorical cross-entropy, left in each panel), the image reconstruction objective (“recon”), and the “combined” classify and reconstruct semi-supervised autoencoder objective. For each comparison between a given neural network layer and brain area, we computed the canonical correlations of the first 10 CCA components, and averaged their values. We repeated this process for 15 random draws of the neural network unit activations, and display the distribution of the resultant CCA correlation values (over those 15 draws) as a box and whisker plot. Lines within the filled bar indicate the mean, and filled rectangle corresponds to the interquartile range.

For the “classify” model, IT was best described by the latent representation (z), whereas V4 was better described by the conv3 layer, which is earlier in the hierarchy. This in line with previous work (e.g., Refs. 4,17) showing that deeper layers of task-trained neural networks are better matches to brain regions deeper in the ventral stream’s visual hierarchy. For contrast, with our “reconstruct” and “combined” objectives – which involve an unsupervised component – the best match to both the V4 and the IT data, was from the latent representation (z) of the neural network. This suggests that the specific alignment of which brain area is best matched by which layer of an artificial neural network model could depend on the task for which the artificial neural network is optimized.

To determine which objective function led to neural networks that best match each brain area, we identified the layer of each network that gave the highest mean canonical correlation with each brain region. For area IT, this was the latent representation (z) in all models; whereas for area V4, this was the latent representation (z) for the “reconstruct” and “combined” models, and layer conv3 for the “classify” model. We then compared these best-layer mean canonical correlation values between neural network models, for each brain area, to determine which model(s) best described the brain data.

For area IT, the “combined” model had the highest mean canonical correlation value (0.265 +/− 0.002: mean +/− standard error, over 15 random samplings of neural network unit activations; see Methods), followed closely by the “reconstruct” model (0.262 +/− 0.003: mean +/− standard error, over 15 random samplings of neural network unit activations), and more distantly by the “classify” model (0.240 +/− 0.005: mean +/− standard error, over 15 random samplings of neural network unit activations). The differences between models was statistically significant in all cases (p = 1×10^−2^ for comparing the “combined” and “reconstruct” models; p = 2×10^−6^ for comparing the “combined” and “classify” models; and p = 2×10^−6^ for comparing the “reconstruct” and “classify” models. All comparisons were done with one-tailed Wilcoxon rank sum tests.)

Our findings in area V4 mirrored those from IT: the “combined” model had the highest mean canonical correlation value (0.245 +/− 0.004: mean +/− standard error, over 15 random samplings of neural network unit activations), followed closely by the “reconstruct” model (0.239 +/− 0.005: mean +/− standard error, over 15 random samplings of neural network unit activations), and more distantly by the “classify” model (0.21 +/− 0.01: mean +/− standard error, over 15 random samplings of neural network unit activations). The differences between models was statistically significant in all cases (p = 2×10^−3^ for comparing the “combined” and “reconstruct” models; p = 2×10^−6^ for comparing the “combined” and “classify” models; and p = 2×10^−6^ for comparing the “reconstruct” and “classify” models. All comparisons done with one-tailed Wilcoxon rank sum test.)

Having identified the best models, and motivated by the analyses by in Ref. 12, we asked how the different attributes in the input images – both categorical and non-categorical -- were represented by the different models. We first tested the position sensitivity of the units in each layer of the neural network model, using test images of clothing items on the natural scene backgrounds (Fig. 3AB; see Methods). For both the “reconstruct” and “combined” models, the position sensitivity increased monotonically with increasing depth. Whereas, for the “classify” model, the position sensitivity decreased between conv4 and the subsequent latent representation (z). (Notably, all layers before the latent representation in our model are convolutional, whereas the latent representation is a *fully connected layer*. For comparison, the authors of Ref. 12 showed position sensitivity in their model – trained purely for categorization – that increased monotonically with depth, for the 6 convolutional layers of their model. This could seem at odds with the fact that our latent representation is less position sensitive than are the previous layers. However, the fully connected nature of this layer will tend to remove position information, and hence we believe that our results are quite consistent with those of Ref. 12. in terms of position information evolving with depth in fully convolutional neural network layers.).

**Fig. 3:**
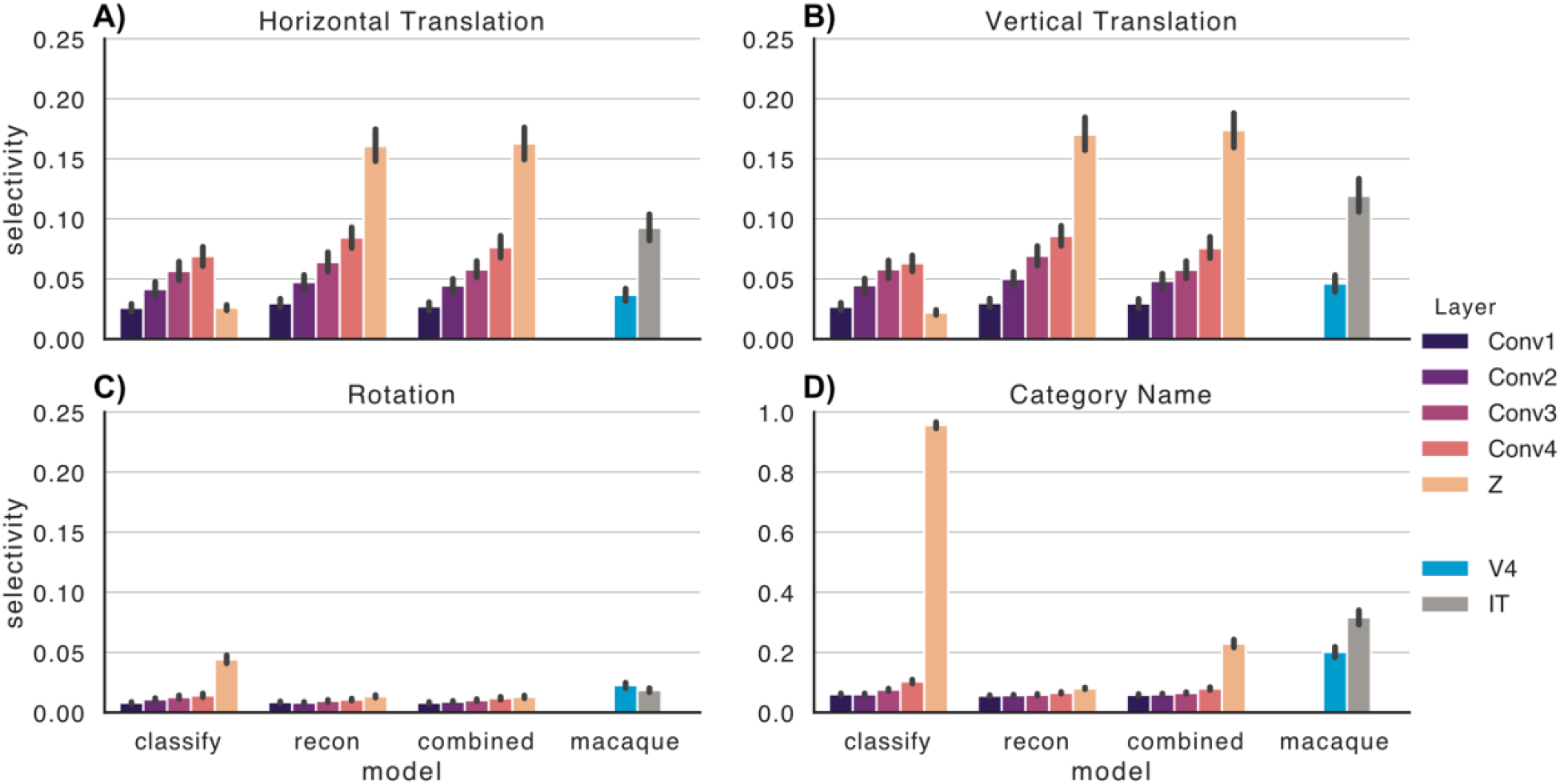
Selectivity for visual scene attributes. Selectivity of units in the fully trained convolutional models optimized under “classify” objective (categorical cross-entropy), “reconstruction” objective, and the “combined” classify+reconstruct semi-supervised autoencoder objective^20^. We measured property selectivity of both categorical (**D**) and continuous valued category-orthogonal properties (**A, B, C**) on units in the multi-electrode array data from Hong et al. (2016), and from units in each layer of the computational model encoders. We defined selectivity for categorical information on each unit in the dataset as the absolute value of that unit’s discriminability (one-vs-all d-prime). We defined selectivity for continuous valued attributes (horizontal and vertical position) on each unit as the absolute value of the Pearson correlation coefficient. Unit activities for models were sampled using 10000 held out test images to generate activations at each layer of the model. We randomly sampled 250 units from each layer of each model for the analysis. Error bars show 95% confidence intervals over the observed set of units.

For comparison, we show the position selectivity from the neurons observed in the monkey experiments, which show increasing position selectivity between V4 and IT.

Next, we tested the rotation selectivity of each of the units in our models. Those were quite low for the units in all of the models, as they were in V4 and IT of the monkey (Fig. 3C). The one exception to this is the latent representation of the “classify” model, which stood out for its high rotation selectivity. Finally, we assessed the category selectivity of the units in each of our models, and show them alongside the corresponding data from monkey V4 and IT (Fig. 3D). Notably, the latent space of the “classify” network stands out for its high category selectivity, compared with the other network models, and the monkey data.

Importantly, the monkey data in Fig. 3 were derived from the images shown to the monkeys, whereas we computed the selectivities of our neural network model units on the images of clothing items superimposed on nature image backgrounds. We did this because the image categories in those images (clothing images) match those on which the network models were trained; these are different from the objects in the images shown to the monkeys. This is a potential limitation in the comparisons between network models and monkey data in Fig. 3.

## Discussion

Here, we studied a supervised learning model (trained to classify objects in images), an unsupervised learning model (trained as an autoencoder to generate compressed representations of input images that suffice for their reconstruction), and a semi-supervised model (trained to both classify objects and enable image reconstruction from its latent representation). We asked which objective function led to neural network models whose image representations most closely match those observed in the ventral stream of the primate visual cortex, and found that the best match was the semi-supervised model. The unsupervised model was close behind, while the supervised model lagged more substantially behind the other two. This suggests that accurate descriptions of ventral stream computations should involve unsupervised learning objectives (e.g., image reconstruction). We also characterized the depth-depending evolution of categorical and non-categorical information in these models, with an aim towards understanding how the different objectives affect the representation of different image attributes at different depths in the neural networks.

We are not the first to explore unsupervised learning algorithms as models of ventral stream (VS) computation. For example, the classic “sparse coding” models showed that unsupervised autoencoders formed image representations similar in many ways to those observed in primary visual cortex (V1)^18,24,25^. More recent work showed that better descriptions of primate V1 responses could be obtained with supervised learning algorithms trained for object recognition ^16,17^ than with the unsupervised algorithms^16^, or with wavelet bases that mimic those learned by the unsupervised learning algorithms^17^. Those works did not look at deeper areas of the VS (e.g., V4 or IT), nor did they study the different objectives in the same neural network architectures.

Two concurrent studies^14,15^ overcome these challenges – as does this paper. Those studies also investigated unsupervised deep learning algorithms, and found that they better matched VS image representations than do supervised algorithms. This is at odds with earlier studies (e.g. Refs. 2,4), which suggested that supervised algorithms (like our “classify” model) would be the best, although it is in-line with other work that questioned whether “pure” object recognition systems really were the best models of ventral stream physiology^26,27^. To this body of work, we add the observation that semi-supervised algorithms (inspired by the machine learning work of Ref. 20) could be even better than the “pure” unsupervised learning algorithms.

Compellingly, and in line with our findings, recent studies of human perceptual judgments of object categories showed that neural networks that combined an image-generative component with a classification component, gave closer matches to the human behavioral data than did networks without the generative component^28^. In other words, both in terms of human perceptual judgments^28^, and primate neurophysiology (this work), our best understanding of VS computation might be in terms of a combination of different task objectives, that include object recognition and image reconstruction. I.e., semi-supervised models might form our best models of the VS.

Somewhat surprisingly, we found that categorization performance in our “combined” model was nearly as good as in our “classify” model (Fig. 1D), even though the units in the “combined” model were overall less category-selective than were the units in the “classify” model (Fig. 3D). This apparent contradiction is explained by a recent machine learning study^29^, which trained neural networks for object categorization, using regularization that penalized category selectivity in all but the readout layer. This led to networks with much lower single-unit category selectivity, but no commensurate loss in categorization performance at the read-out stage. Thus, the link between single-unit category selectivity, and overall network categorization performance, is surprisingly weak.

Importantly, our goal here was not necessarily to obtain state-of-the-art models of the primate VS. Rather, it was to compare different objective functions within the same architecture, to see which was a better match to the VS. Some recent work of ours^16^ does push more towards obtaining state-of-the-art models, and finds that networks trained end-to-end to predict V1 firing rates achieve higher performance than is obtained using regression against the unit activations from VGG-16 (a pre-trained object classification network). That suggests that there is something more going on in primate VS than “just” object recognition, although another study concurrent to that one^17^ found that regression on VGG-16 activations was slightly better than end-to-end trained models. For many reasons (different datasets, and different inclusion criteria for neurons, for example), direct comparison of performance measures between those studies is difficult. As such, an important future area of work is to systematically sample the space of architectures and objective functions, to find the best one. Our work suggests that semi-supervised objectives are strong candidates for that work, and we are encouraged by efforts like the Brain-Score platform^30^, to facilitate quantitative comparison between models.

One natural question that arises is about our decision to train our models on images of fashion items superimposed on natural image backgrounds, as opposed to other datasets (e.g., ImageNet). We chose this approach because it yielded images of naturalistic objects (clothing items) with rich natural image backgrounds, yet was parametric in the location and orientation of the objects, and highly tractable computationally. The same is not true of ImageNet or other “typical” computer vision benchmark tasks. Moreover, being able to procedurally generate new examples (of clothing items on nature image backgrounds) during training gave effectively endless variation in the training data that improved the training of our models.

Moreover, while we chose canonical correlation analysis (CCA) for comparing neural data to neural network models, many recent studies^2,4,14–17^ (including some of our own^16,31^) used instead analyses based on representational dissimilarity matrices (RDM), or regression between neural network unit activations and recording neuronal activities. While we like the RDM and regression approaches, all of them (including CCA) have important limitations, leaving it unclear which is the best method to compare neural networks to brains. First, RDM compares matrices of image-by-image (or category-by-category) dissimilarity in activation vectors in the neural network, to those obtained from the brain^32^. In this approach, even if the neurons in the brain were exactly recapitulated by units in the neural network, the RDM analysis could still show a poor match if there are *other* units in the neural network that do not match those in the brain from which the experimenters recorded. Given that neural data is invariably subsampled (not all neurons are recorded), this can be serious limitation. Regression-based approaches get around this challenge by attempted to reconstruct the neuronal activities from the neural network unit activations. A downside to this approach is the need for heavy regularization to prevent overfitting, and the difficulty in deciding how to average the prediction quality (usually a correlation, or fraction of explained variance) over neurons to get ensemble statistics. Those values are typically just averaged over cells, but neurons’ activations are usually correlated with each other, so that averaging can be problematic. CCA attempts to circumvent these issues, by finding linear combinations of neural network unit activations, that most correlate with linear combinations of neuronal activities. When multiple components are obtained, they are each independent of one another, enabling us to average over their correlation values (we used 10 CCA components in this study). For these reasons and others, an increasing number of neuroscientists are using CCA for analyses like the one presented here^22,23^. We do not intend here to argue that any one of these methods is better than any other. All of them have limitations, and an important avenue for research is to determine, on principled grounds, which approach is best for different types of comparisons between brains and artificial neural networks.

It is important to mention that this study had several important limitations. First, we studied only a single neural network architecture. In principle, different results could be obtained with other architectures. At the same time, the concurrent results from other groups^14,15^ (using other architectures and image datasets), showing that unsupervised learning provides better VS models than does supervised learning, increases our confidence in our findings. Second, our results from images of fashion items on nature scene backgrounds could, in principle, fail to generalize to other settings. On the other hand, natural images have strong statistical regularities^33,34^, suggesting that, so long as one samples broadly from the realm of realistic images, the specific images chosen may not be overly important. Our images – of real-world objects on nature image backgrounds – should thus not pose any serious issues.

We conclude by noting that a key open question in neuroscience is to find the computational objectives that describe the visual ventral stream. Our work suggests that semi-supervised objectives, combining object recognition with scene reconstruction, may be promising candidates.

## Materials and Methods

### Primate Electrophysiology

Neural recordings were originally collected by the DiCarlo lab (Ref. 12) and shared with us for this analysis. In brief, neural recordings were collected from the visual cortex of two awake and behaving rhesus macaques using multi-electrode array electrophysiology recording systems (BlackRock Microsystems). Animals were presented with a series of images showing 64 distinct objects from 8 classes rendered at varying position in the animal’s visual field, and with variation rotations. After spike-sorting and quality control this resulted in well-isolated single units from both IT (n=168) and V4 (n=128); higher-order areas in primate visual cortex. A full description of the data and experimental methods is given by Ref. 12.

### Dataset and Augmentation

Our goal was to study the object representations, scene reconstruction, and representation of non-categorical information, within artificial neural networks. To achieve that goal, we trained the neural networks to take in images, and either categorize the objects within them, reconstruct the images, or categorize the objects *and* reconstruct the input (i.e., a semi-supervised autoencoder^20^). To train these networks, we required images that varied in categorical, and in non-categorical, properties. For that reason, we constructed images of clothing items superimposed at random locations over natural image backgrounds.

To achieve this goal, we used all 70,000 images from the Fashion MNIST dataset, a computer vision object recognition dataset comprised of images of clothing articles from 10 different categories. We augmented this dataset by superimposing those 28×28 pixel images onto 112×112 pixel frames, with the center locations drawn randomly from a uniform distribution spanning 75% of the image field. Images were shifted according those randomly drawn dx and dy values, and rotated according to randomly drawn angles between −54 and +54 degrees. After applying positional and rotational shifts, the objects were superimposed over random patches extracted from natural images from the BSDS500 natural image dataset to produce simplified natural scenes which contain categorical (1 of 10 clothing categories) and non-categorical (position and rotation shifts) variation. Random 112×112 pixel patches from the BSDS500 dataset were gray scaled before the shifted object images were added to the background patch (Fig 1A). All augmentation was performed on-line during training. That is, every position shift, rotation shift, and natural image patch was drawn randomly every training batch instead of pre-computing shifts and backgrounds. This allows every training batch to be composed of unique combinations of objects, backgrounds, rotations, and shifts, helping to prevent overfitting. This approach yielded 112×112 pixel images that contained the clothing item, at a random location and orientation, with a nature image background.

### Computational models

The convolutional models were constructed by sequentially combining convolutional layers, followed by an all-to-all connected layer (z). Each convolutional layer receives as input a spatially arranged map from the prior layer. A filter kernel is multiplied against the input at each spatial location in the input, and the resultant value is added to the bias and passed through the nonlinear activation function.

The models described in our paper were constructed according to the table below. The first 4 layers were convolutional, whereas the latent layer (z) was densely connected.

|  | *Output Size* | *Kernel*<br>*Size* | *Activation*<br>*Function* | *Dropout*<br>*rate* | *Batch Normalization*<br>*Momentum* |
| --- | --- | --- | --- | --- | --- |
| *Input* | 112 x 112 | N/A | N/A | N/A | N/A |
| *Layer 1* | 56x56x16 | 3x3 | LeakyReLU | 25% | 0.8 |
| *Layer 2* | 28x28x32 | 3x3 | LeakyReLU | 25% | 0.8 |
| *Layer 3* | 14x14x64 | 3x3 | LeakyReLU | 25% | 0.8 |
| *Layer 4* | 7x7x128 | 3x3 | LeakyReLU | 25% | 0.8 |
| *Latent, z* | 500 |  | Linear | 0% | 0.8 |

Models using the “reconstruct” objective, and the “composite” classify-and-reconstruct objective (see below) need an additional generator network to reconstruct the original stimulus input from the latent representation. The generator network (G) uses a residual convolutional neural network (ResNet) which has achieved state of the art performance in natural image generation. The generator network uses is comprised of deconvolutional layers and its architectural hyperparameters directly mirror those in the convolutional encoder. We chose this generator network structure because it led to better performance (lower sums of squared errors in image reconstruction) than other generators we had tried, including ones that mirrored the encoding side of our network models. We do not claim that this generator model describes anything about the biology: it is there instead to enable an image to be decoded from the latent representation, to help test whether the latent representation contains sufficient information for that reconstruction.

Our models can be found on Github (https://github.com/elijahc/vae).

### Objective functions and training parameters

Models optimized for classification use categorical cross-entropy for the objective function. Categorical cross-entropy (XENT) is a commonly used objective function in machine learning to train neural network classifiers. Multilabel cross-entropy is calculated according to the equation below where M is the total number of classes

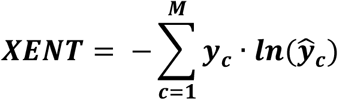

Here, ***y***_***c***_ is the true category label, represented as a one-hot vector, and ***ŷ*** _***c***_ is the network output obtained from the linear readout of the latent state (see Fig. 1).

Models optimized for reconstructing the original input scene use pixel-wise sum of squared error (SSE) between the input and the generator’s output 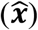.

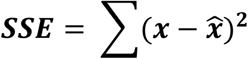

Models optimized for both objectives (i.e., the “combined” objective) were optimized for the sum of the two: their objective function was *SSE + XENT*.

Notably, other objective functions could also have been used for the reconstruction loss, in place of our SSE objective. One example would be the contrastive loss (as in Ref. 14). We do not claim that the SSE is the only (or even the “best”) loss function for the unsupervised learning component. Minimizing this loss does, however, force the network’s latent representation to retain sufficient information about the input to enable its reconstruction.

We trained each model in our experiment until classification accuracy plateaued on a validation dataset of 512 objects from the 10,000 test images in the fashion MNIST dataset.

### Model Evaluation

#### Canonical Correlation Analysis (Fig. 2)

We quantified the similarity of each models’ layer-wise selectivity to corresponding layers in primate ventral stream using Canonical Correlation Analysis (CCA)^22^. CCA finds a set of weights used to project both the primate electrophysiology results and our own model unit activations into a lower dimensional space and measures the correlation of the projections in this space. The projection weights are optimized to maximize correlation in the lower dimension. We use 10 projection dimension for this analysis and report the average over the (optimized) correlations of those 10 dimensions. In analogy to the monkey experiments, we performed these analyses on randomly-chosen sets of 250 units from our models; this approximates the number of pseudo-randomly sampled of neurons with the implanted electrode arrays. While these 250 units represent 50% of our latent space (z), the fraction of neurons sampled from monkey V4 or IT in the physiology experiments was much lower.

We repeated the analysis for 15 different random draws of unit activations and report the distribution of correlations over those 15 draws (Fig 2).

#### Feature Selectivity (Fig. 3)

After training performance plateaus, 5-fold sampling of 250 randomly chosen unit activations from each layer in the encoder model (Fig 1B) were used in comparisons with primate ventral stream electrophysiology. Unit activations were generated using a random sample from held out test images (not used during training). As in a (simulated) electrophysiology experiment, each image was input to the network, and the corresponding unit activations were recorded. We then analyzed these unit activations in the same way as we did the firing rates recorded in monkey visual cortex, described below.

First, we measured selectivity of our artificial neurons to different image attributes, in the same way as Ref. 12 (they call these measures “performance” instead of selectivity). For continuous-valued scene attributes (e.g. horizontal position) we measured selectivity as the absolute value of the Pearson correlation between the neuron’s response and that attribute in the stimulus image. For categorical properties (e.g. object class) we measure selectivity as the one-vs-all discriminability (d’).

## Acknowledgements

We would like to thank the DiCarlo lab for sharing their primate electrophysiology recordings with us. Special thanks to Alon Poleg-Polsky for thoughtful discussion and direction and to Doug Crawford, Shaiyan Keshvari, Martin Schrimpf, Rachel Sewell, and Heidi Sjoberg for helpful feedback on the manuscript. EC acknowledges funding by an NDSEG fellowship through the US Department of Defense. JZ acknowledges funding from CIFAR, the A.P. Sloan Foundation, Google, the Canada Research Chairs Program, and the Natural Sciences and Engineering Research Council of Canada (NSERC, RGPIN-2019-06379).

